# Calcium ions trigger the exposure of phosphatidylserine on the surface of necrotic cells

**DOI:** 10.1101/2020.08.21.260695

**Authors:** Yoshitaka Furuta, Omar Pena Ramos, Zao Li, Zheng Zhou

**Affiliations:** Verna and Marrs McLean Department of Biochemistry and Molecular Biology, Baylor College of Medicine, Houston, TX 77030; School of Pharmacy, Kanazawa University, Kakuma-machi, Kanazawa, Ishikawa, 920-1192, Japan; Memory & Brain Research Center, Department of Neuroscience, Baylor College of Medicine, Houston, TX 77030; Genentech Inc., South San Francisco, California

## Abstract

Intracellular Ca^2+^ level is under strict regulation through calcium channels and intracellular Ca^2+^ storage pools such as the endoplasmic reticulum (ER). Mutations in certain ion channel subunits, which result in the mis-regulation of Ca^2+^ influx, cause the excitotoxic necrosis of neurons. In the nematode *Caenorhabditis elegans*, six mechanosensory (touch) neurons are induced to undergo excitotoxic necrosis by dominant mutations in the DEG/ENaC sodium channel subunits. These necrotic neurons are subsequently engulfed and degraded by neighboring hypodermal cells. We previously reported that the necrotic touch neurons actively expose phosphatidylserine (PS), an “eat-me” signal, to attract engulfing cells. However, the upstream signal that triggers PS externalization remained elusive. Here we report that a robust and transient increase of cytoplasmic Ca^2+^ level occurs prior to the exposure of PS on the surfaces of necrotic neurons. We further found that inhibiting the release of Ca^2+^ from the ER, either pharmacologically or genetically through mutations in the gene encoding calreticulin, the ER Ca^2+^ chaperon, impairs PS exposure on necrotic neurons. On the contrary, inhibiting the re-uptake of cytoplasmic Ca^2+^ into the ER induces ectopic necrosis and PS exposure. These findings indicate that high levels of cytoplasmic Ca^2+^ is necessary and sufficient for PS exposure. Remarkably, we found that PS exposure occurred independently of other necrosis events. On the other hand, apoptotic cells, unlike necrotic cells, do not depend on the ER Ca^2+^ pool for PS exposure. Our findings reveal a necrotic neuron-specific, “two-step Ca^2+^-influx” pathway that promotes PS exposure on cell surfaces. This pathway is initiated by the modest influx of Ca^2+^ from the extracellular space and further boosted by the release of Ca^2+^ from the ER into the cytoplasm.

**Author Summary:** Necrosis is a type of cell death that exhibits distinct morphological features such as cell swelling. Many environmental insults induce cells to undergo necrosis. Necrotic cells expose phosphatidylserine (PS) – a type of phospholipid – on their outer surfaces. Receptor molecules on phagocytes detect phosphatidylserine on necrotic cells and subsequently initiate the engulfment process. As necrosis is associated with stroke, cancer, neurodegenerative diseases, heart diseases, and inflammatory diseases, studying necrotic cell clearance has important medical relevance. In the model organism the nematode *C. elegans*, by utilizing dominant mutations in ion channels that induce neurons to undergo necrosis, we previously identified membrane proteins that promote the exposure of phosphatidylserine on necrotic cell surfaces. Here, using the same experimental system, we further discover that the necrosis insults trigger an increase of the cytoplasmic Ca^2+^ level, which in turn promotes PS externalization in necrotic cells. Furthermore, we find that the Ca^2+^ pool in the endoplasmic reticulum is necessary for the rapid increase of cytoplasmic Ca^2+^ that helps initiate necrosis. This Ca^2+^-regulated event is not observed when cells undergoing apoptosis (a form of cell suicide) expose PS. Our findings reveal a novel upstream regulatory mechanism that promotes necrotic cell clearance in animals.

## Introduction

Cells undergoing necrosis - a type of cell death morphologically distinct from apoptosis - display cell and organelle swelling, excessive intracellular membranes, and the eventual rupture of the intracellular and plasma membranes [1,2]. Necrosis is frequently observed during cell injury, and is closely associated with stroke, heart diseases, inflammatory diseases, diabetes, cancer, and neurodegeneration [3-8]. Although necrosis has historically been considered an uncontrolled cell death event, recent discoveries from multiple organisms demonstrated that cells possess genetic programs that trigger necrosis in response to extracellular stimuli [9-12]. In the nematode *Caenorhabditis elegans*, dominant (*d*) mutations in certain ion channel subunits of the DEG/ENaC (degenerin/epithelial sodium channel) superfamily, in the nicotinic acetylcholine receptor, in trimeric GTPases, and in a few other proteins induce neurons of specific identities to undergo a type of necrosis that resembles the excitotoxic necrosis of mammalian neurons [10]. One such gene is *mec-4*, which encodes a core subunit of a multimeric, mechanically gated sodium channel that belongs to the DEG/ENaC family and functions in the mechanosensory (touch) neurons for sensing gentle touch along the worm body [13]. Dominant mutations in *mec-4* trigger the necrosis of all the six touch neurons (AVM, PVM, ALML/R and PLML/R) [13,14]. Unlike apoptosis, the necrosis triggered by the *mec-4(d)* mutations does not require the function of the CED-3 caspase [15]. Furthermore, many of the cellular events occurring during necrosis, including those leading to cellular elimination, are different from those occurring during apoptosis [1,2].

Cells undergoing necrosis, like those undergoing apoptosis, are recognized, engulfed, and degraded by neighboring engulfing cells [16,17]. Apoptotic cells are known to present phosphatidylserine (PS), a membrane phospholipid, on their outer surfaces to attract the phagocytic receptors on engulfing cells [18]. PS is thus referred to as an “eat me” signal that is recognized by engulfing cells, triggering subsequent engulfment [18]. We previously discovered that the outer surfaces of necrotic touch neurons in *mec-4(d)* mutant worms expose PS, like apoptotic cells [19,20]. In turn, this PS interacts with the phagocytic receptor CED-1 resides on the surfaces of neighboring engulfing cells, allowing necrotic cells to be recognized by engulfing cells [19]. We found that PS was actively exposed on the surface of necrotic neurons while the plasma membrane remained intact. In addition, we identified two proteins that act in parallel to promote the externalization of PS from the inner plasma membrane leaflet to the outer leaflet during necrosis [19]. One of these proteins is CED-7, the *C. elegans* homologs of the mammalian ABCA transporter [19,21]. Mammalian ABCA possesses the PS-externalization activity [22], and CED-7 is essential for promoting PS exposure on the outer surfaces of apoptotic cells [20,23]. The other protein is ANOH-1, the *C. elegans* homolog of mammalian phospholipid scramblase TMEM16F, which catalyzes the random, bi-directional “flip-flop” of phospholipids across the membrane bilayer [19,24]. Whereas CED-7 is expressed broadly and its function is needed in both apoptotic and necrotic cells for efficient PS exposure, ANOH-1 is specifically expressed in neurons and specifically acts in necrotic neurons, but not apoptotic cells, to facilitate PS exposure [19-21]. Mammalian TMEM16F was reported to promote PS exposure on the surfaces of platelets during the blood clotting process [24-26]. To our knowledge, *C. elegans* ANOH-1 is the first phospholipid scramblase reported to promote PS exposure on necrotic cells.

The proteins responsible for PS externalization on *C. elegans* touch neurons during necrosis are presumably activated by upstream signal(s). However, the identity of such upstream signal(s) remained unknown. In comparison, during apoptosis, caspases are known to be the critical upstream molecules that trigger the exposure of PS and other cellular events. In living cells, PS is almost exclusively localized to the inner leaflet of the plasma membrane, at least partially due to the ATP-dependent aminophospholipid translocase activity that selectively returns PS from the outer to the inner leaflet [18]. In mammalian cells undergoing apoptosis, caspase 3, an effector caspase that is activated by upstream apoptosis signals, cleaves and inactivates the aminophospholipid translocase ATP11C [27]. Caspase 3 also cleaves Xk-related protein 8 (Xk8), another phospholipid scramblase, and this cleavage results in the activation of Xk8’s scramblase activity [28]. By inactivating the translocase ATP11C and activating the scramblase Xk8, caspase cleavage enables the externalization of PS on the surface of apoptotic cells [27-29]. Similarly, evidence suggests that CED-8, the *C. elegans* homolog of mammalian Xk8, is cleaved and activated by CED-3 during apoptosis to facilitate PS exposure [28,29].

Caspase activity has not been found to regulate excitotoxic necrosis, including the necrosis of the touch neurons triggered by *mec-4(d)* mutations [15,30,31]. Rather, intracellular calcium ions were considered important signaling molecules that induce the necrosis of multiple types of cells [32-34], including the *C. elegans* touch neurons [35]. The *mec-4(d)* mutations alter the conformation of the mechanosensory Na^+^ channel (of which MEC-4 is a subunit) and make it permeable to extracellular Ca^2+^ [36]. Remarkably, *mec-4(d)* induced necrosis is partially suppressed by loss-of-function (*lf*) mutations in *crt-1* – which encodes calreticulin, the ER residential Ca^2+^ chaperone that facilitates the accumulation of Ca^2+^ in the ER – and by inactivating genes encoding the ER Ca^2+^-release channels [35]. Additionally, *mec-4(d)*-induced necrosis is suppressed by chemical treatment that impairs the release of Ca^2+^ from the ER to the cytoplasm [35]. Based on the knowledge that (1) the cytoplasmic Ca^2+^ level is generally much lower than that in the extracellular space or intracellular Ca^2+^ pools [37], (2) that the ER is one of the primary storage pools for intracellular Ca^2+^ [37], and (3) that the Ca^2+^ release channels in the ER are activated by the increase of cytoplasmic Ca^2+^ [38], Xu et al [35] proposed that the small amount of extracellular Ca^2+^ that entered the touch neurons through the MEC-4(d)- containing Na^+^ channel further induced a robust Ca^2+^ release from the ER to the cytoplasm. Moreover, they propose that necrosis is induced once the level of cytoplasmic Ca^2+^ reaches a certain threshold [35]. Closely related to this discovery and to our investigation of PS exposure on necrotic cells, the scramblase activity of mammalian TMEM16F is known to be dependent on Ca^2+^ [24].

We set out to investigate whether cytoplasmic Ca^2+^ triggers PS exposure on necrotic cells and whether the ER, as an intracellular Ca^2+^ pool, takes part in the externalization of PS. Using a fluorescently tagged-MFG-E8, a secreted reporter that detects extracellular PS [19,20], we quantitatively monitored PS exposure on the cell surface. Previously, the level of Ca^2+^ in touch neurons has not been monitored over the course of necrosis. To study the relationship between cytoplasmic Ca^2+^ and PS exposure, we developed a touch neuron-specific Ca^2+^ indicator. By monitoring the Ca^2+^ indicator and the PS reporter, we observed a close relationship between a surge of cytoplasmic Ca^2+^ and the exposure of PS on the cell surface of necrotic cells. We have further observed that the release of Ca^2+^ from the ER is essential for the exposure of PS, that the exposure of PS is regulated by Ca^2+^, and that this phenomenon operates independent of other necrosis events. Our findings reveal a necrotic cell-specific, Ca^2+^-dependent regulatory mechanism that is facilitated by the ER and that acts to trigger the exposure of the “eat me” signal and the clearance of necrotic cells.

## Results

### Multiple kinds of necrotic neurons expose phosphatidylserine (PS) on their surfaces

The six *C. elegans* touch neurons, when undergoing necrosis in the *mec-4(e1611)* dominant mutant strain, expose PS on their surfaces. This PS signal is detected by a mCherry-tagged MFG-E8, a PS-binding protein that is expressed in and secreted out of neighboring hypodermal cells (**Fig 1 c-d**) [19]. Conversely, MFG-E8::mCherry does not detect PS on the surfaces of live touch neurons (marked by P_*mec-7*_GFP) in wild-type larvae (**Fig 1 a-b**) [19]. To determine whether PS exposure is a common feature for necrotic neurons of different kinds and induced to die by different insults, we examined a number of necrotic neurons in *deg-1(u38), trp-4(ot337), unc-8(n491sd)*, and *deg-3(u662)* dominant mutant L1 larvae. *deg-1* encodes a DEG/ENaC family sodium channel subunit [39]. *trp-4* encodes a transient receptor potential (TRP) channel expressed in dopaminergic neurons and a number of other neurons [40]. *unc-8* encodes another DEG/ENaC family sodium channel subunit that functions as a critical regulator of locomotion [41,42]. *deg-3* encodes a ligand-gated calcium channel belonging to the nicotinic acetylcholine receptor family [43]. Dominant mutations in these genes induces necrosis of neurons: the *deg-1(u38)* mutation causes necrosis of the IL1 sensory neurons and the AVD, AVG, and PVC interneurons [44]; the *trp-4(ot337)* mutation results in the necrosis of dopaminergic neurons and a few non-dopaminergic neurons, including two DVA and DVC, two mechanosensory neurons in the tail [31]; the *unc-8(n491sd)* mutation induces the necrosis of a number of motor neurons at L1 stage [45]; and the *deg-3(u662)* mutation causes the necrosis of a number of sensory neurons including the touch neurons and IL1 and PVD, and interneurons AVG and PVC [43]. *deg-1(u38), trp-4(ot337), unc-8(n491sd)*, and *deg-3(u662)* mutants all exhibited strong PS signal on the surfaces of necrotic cells **(Fig 1 e-l)**, indicating that PS exposure is a common event occurring on necrotic sensory neurons, interneurons, and motor neurons of different identities and induced to undergo necrosis by mutations of different genes.

**Figure 1.**
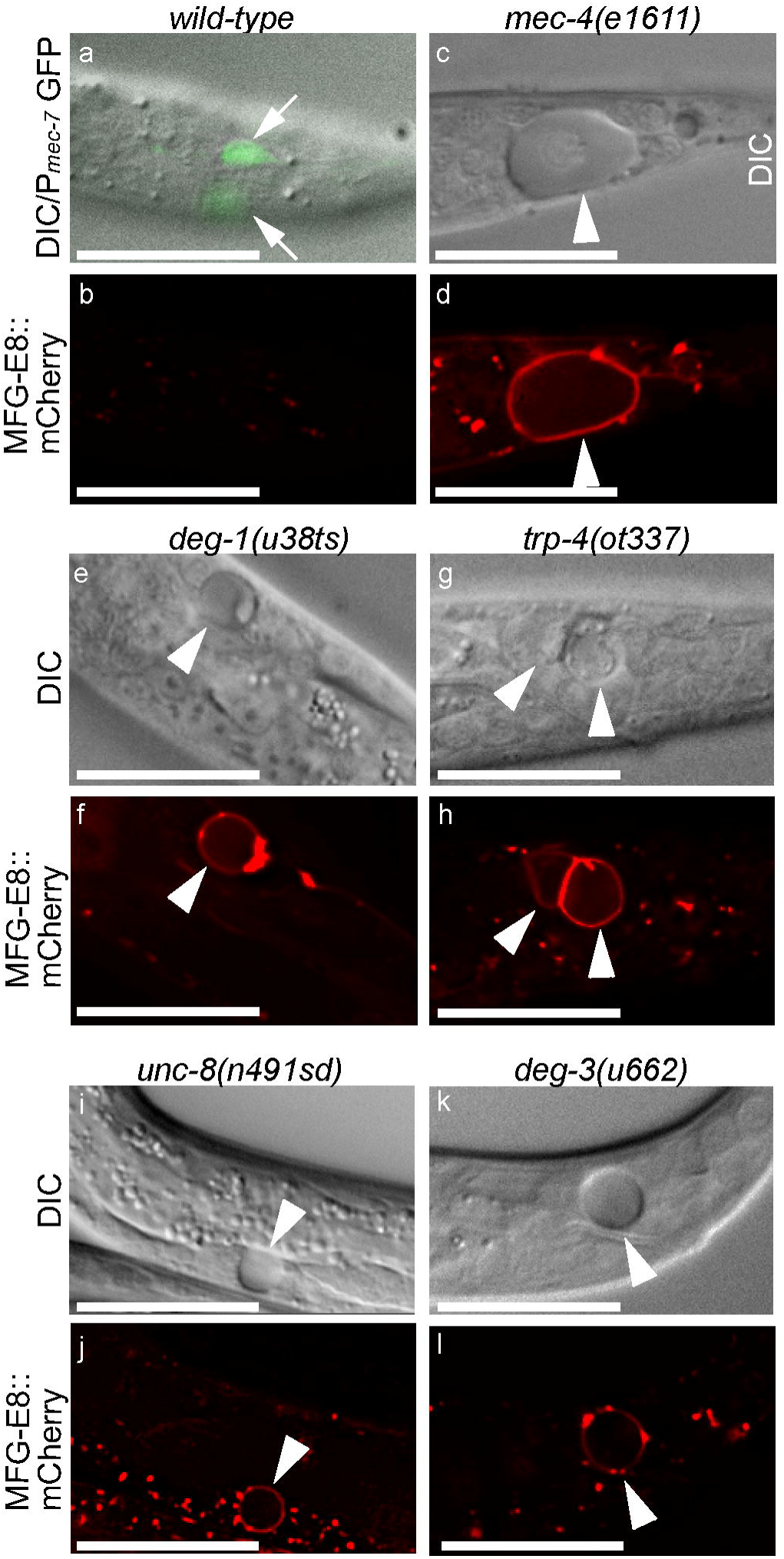
PS exposure is a common phenomenon observed on various necrotic neurons. DIC (a, c, e, g, i, k) and epifluorescence (b, d, f, h, j, l) images of two live touch neurons (white arrows) and six necrotic neurons (white arrowheads) in the L1 larvae of wild-type (a-b) and different mutant strains expressing *P*_*dyn-1*_*mfg-e8::mCherry*, including the PLM touch neuron in the tail (a-d), the AVG interneuron in the head (e-f), the DVA and DVC neurons in the tail (g-h), a ventral motor neuron (i-j), and a putative PVD neuron in the tail (k-l). In (a), P_*mec-7*_GFP labels two PLM neurons (white arrows). For (e-f), AVG interneurons in the head are observed at an incubation temperature of 25°C, as *deg-1(u38ts)* is a temperature sensitive mutant. Scale bars are 15*µ*m.

### A robust and transient increase in the cytoplasmic Ca^2+^ level is detected preceding PS exposure on necrotic neurons

To determine whether cytoplasmic Ca^2+^ level changes as predicted during the necrosis of touch neurons [35], we constructed a Ca^2+^ indicator (GCaMP5G) that is expressed in touch neurons under the control of the P_*mec-7*_ promoter [46] and introduced P_*mec-7*_ *GCaMP5G* into the *mec-4(e1611)* mutant strain (**Materials & Methods**). GCaMP5G generates a bright GFP signal only in the presence of Ca^2+^ due to the conformation change of this fusion protein induced by its interaction with Ca^2+^ [47]. The P_*mec-7*_ *GCaMP5G* reporter is able to detect cytoplasmic Ca^2+^; also, because GCaMP5G does not have any ER signal sequence, it does not detect the Ca^2+^ in the ER lumen. We co-expressed GCaMP5G with the PS reporter MFG-E8::mCherry and established a time-lapse recording protocol that allowed us to simultaneously monitor GCaMP5G and MFG-E8::mCherry signals throughout embryonic development. In *mec-4(e1611)* mutants, touch neurons PLML and PLMR (**Fig 2A**) undergo necrosis during late embryonic developmental stage [19]. We monitored the Ca^2+^ and PS signals throughout the necrosis process of PLML and PLMR (**Materials and Methods**) and detected a robust and transient increase of GCaMP5G intensity in the cytoplasm of the PLM neurons (**Fig 2 B-D**). The spike in GCaMP5G intensity occurs prior to cell swelling and PS exposure, indicating that the rise in cytoplasmic Ca^2+^ level precedes these events **(Fig 2 B-D)**. The transient peak in Ca^2+^ subsequently gradually reduces to the basal level (**Fig 2 B-C)**. The recording of multiple samples revealed that although there was certain degree of variation of both the peak Ca^2+^ signal value and the width of the peak, a general pattern of a cytoplasmic Ca^2+^ peak preceding cell swelling and PS exposure remained consistent among necrotic neurons **(Figs 2 (B-C), 4(A-B), and S1C)**.

**Figure 2.**
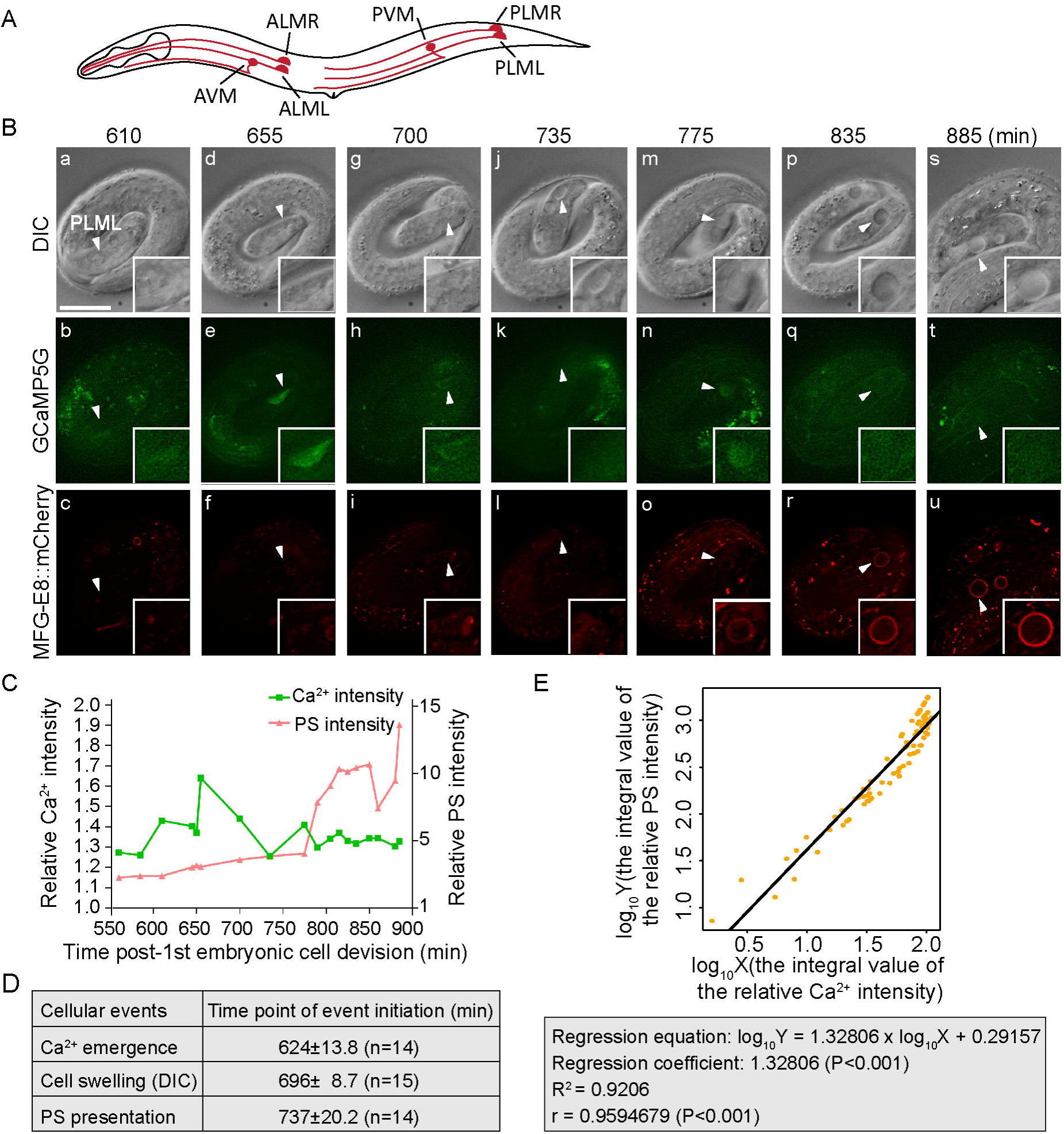
Ca^2+^ signal in the cytoplasm increases prior to the onset of necrosis morphology and the exposure of PS, and displays a linear relationship with the PS level. (A-C) Results of time-lapse recording experiments monitoring cytoplasmic Ca^2+^ levels, cell swelling, and PS signal in or on the surface of a PLML neuron, respectively, are reported here. The strain is homozygous for the *mec-4(e1611)* allele and carries a transgenic array co-expressing P_*mec-7*_GCaMP5G and P_*dyn-1*_*mfg-e8::mCherry*. (A) Diagram of a hermaphrodite indicating the positions and names of the six touch neurons. (B) Time-lapse images following the necrosis of one PLML neuron (white arrowheads). Time points are marked as min post-first embryonic cell division. Recording started at 100 min after the larva reached 2-fold stage (560 min) and ended at 900 min. The GCaMP5G signal appears in (b) and reaches its peak value in (e). PLML starts swelling in (g) and continues swelling in (j,m,p,s). The MFG-E8::mCherry signal on the surface of PLML becomes visible in (o) and increases over time (r, u). The scale bar is 10*µ*m. (C) Relative signal levels (in comparison to the background levels) of GCaMP5G in the cytoplasm, and of MFG-E8::GFP on the membrane surface, of the PLML shown in (A). (D) Summary of the time points when each of the three cellular events initiates in PML neurons during the onset of necrosis. The appearance of Ca^2+^ in the cytoplasm (GCaMP5G), the distinct swelling morphology observed under DIC microscope, and the appearance of PS on the surface (MFG-E8::mCherry). Data represent mean ± standard deviation. n, the numbers of embryos analyzed. (E) Linear regression analysis between the log-transformed relative cytoplasmic Ca^2+^ level in and relative PS level on the surfaces of the PML neurons. The integral values of the Ca^2+^ and PS signals throughout the observed time period (560 min to 900 min post-1^st^ embryonic division) were calculated. Linear regression and Pearson’s correlation coefficient analysis were performed by the R Software. Each plot was derived from time points from 6 independent time-lapse recording series. The data include 75 time points. X: the integral value of the relative Ca^2+^ intensity; Y: the integral value of the relative PS intensity; R^2^: coefficient of determination; r: correlation coefficient.

On the contrary, in live PLM neurons in the *mec-4(+)* embryos, only weak, background levels of GCaMP5G signal were observed (**Fig S1(A-B)**). Although the GCaMP5G signal also fluctuates, both the peak and basal values are much lower than those measured in the necrotic PLM neurons in the *mec-4(e1611)* embryos. In addition, we calculated the integral Ca^2+^ signal levels over the period of 560-800 min post-1^st^ embryonic division and found that in *mec-4(+)* embryos, the mean value in living PLM neurons was approximately 40% of that in necrotic PLM neurons in *mec-4(e1611)* mutants (**Fig S1C(c)**). The low peak value and low integral value of the cytoplasmic Ca^2+^ signal both correlate with the lack of necrosis and PS exposure in live PLM neurons in the *mec-4(+)* embryos.

The above observations demonstrate that the cytoplasmic Ca^2+^ exhibits a unique and dynamic pattern of fluctuation in necrotic neurons that is not observed in their healthy counterparts.

To examine the potential correlation between the signal levels of cytoplasmic Ca^2+^ and cell surface PS, we quantified the integral values of the relative signal intensities over time. We subsequently conducted the linear-regression analysis between the log-transformed integral values of the cytoplasmic Ca^2+^ level and PS level. Remarkably, we identified a very strong positive correlation between these two values (the correlation coefficient r is about 0.96) (**Fig 2E**). Together, the above observations and the analysis of these findings indicate that the cytoplasmic Ca^2+^ makes a strong contribution to the level of PS exposure on the surfaces of necrotic neurons.

### Suppression of the release of Ca^2+^ from the ER by dantrolene represses the exposure of PS on necrotic neurons

To determine whether the exposure of PS on necrotic touch neurons requires the release of Ca^2+^ from the ER, we examined whether blocking the release of Ca^2+^ from the ER in the *mec-4(e1611d)* mutants would affect PS exposure on necrotic neurons. The ryanodine receptor (RyR) and InsP_3_ receptor (InsP_3_R) are two major kinds of Ca^2+^ release channels on the ER membrane [38]. Dantrolene, a small molecular antagonist of ryanodine receptors [48], was reported to suppress *mec-4(d)* induced necrosis in touch neurons [35]. We placed *mec-4(e1611)* mutant hermaphrodites on NGM plates containing different doses of dantrolene (**Materials and Methods**), and measured the cell surface PS intensity on the PLML and PLMR touch neurons in their progeny as newly hatched L1 larvae using MFG-E8::mCherry (**Fig 3**).

**Figure 3.**
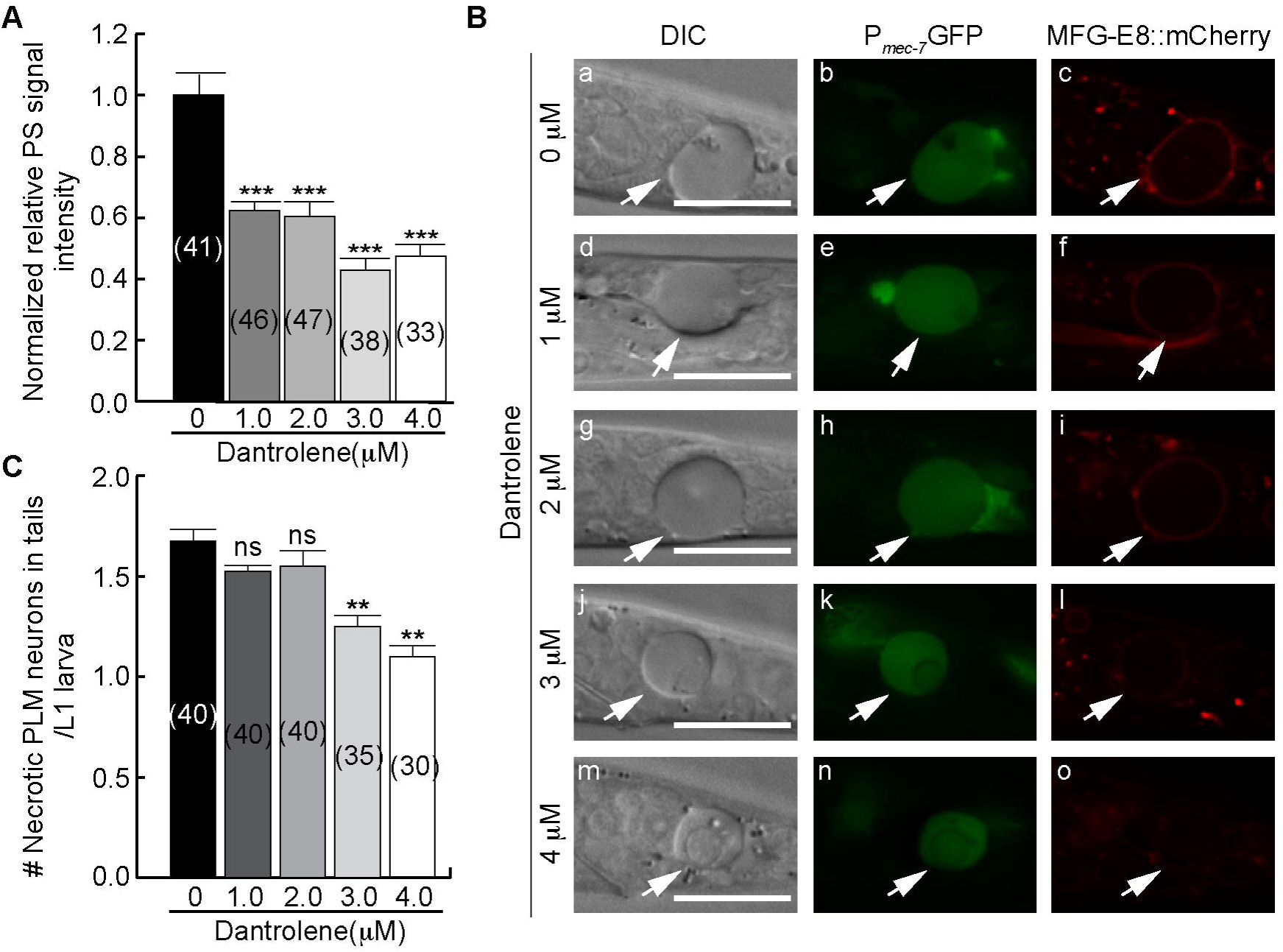
Dantrolene inhibits the exposure of PS on the surface of necrotic neurons. (A) The normalized relative PS intensity (normalized PS_R_) of necrotic PLMs in the tail of newly hatched L1 larvae. Adult worms were placed on NGM plates with different doses of dantrolene and allowed to lay eggs; shortly after hatching, larvae were scored. The “0 μM” samples were given DMSO with no dissolved dantrolene. Bars represent the mean values. Error bars represent standard error of the mean (s.e.m.). Numbers of necrotic PLM neurons scored for each dantrolene concentration are in paratheses. Student *t*-test was used for data analysis: ***, p < 0.001. (B) DIC (a, d, g, j, m), P_*mec-7*_GFP (b, e, h, k, n), and MFG-E8::mCherry (c, f, i, l, o) images of the necrotic PLM neurons (arrows) in the newly hatched L1 larvae. The mother worms were incubated with different doses of dantrolene as labeled. PLM neurons are labeled with P_*mec-7*_GFP. Scale bars are 15 *µ*m. (C) The mean numbers of necrotic PLM neurons in the tails of newly hatched L1 larvae generated by adult worms incubated with different doses of dantrolene. Bars represent the mean values. Error bars represent s.e.m. The numbers of L1 larvae scored after dantrolene treatments are in parentheses. Student *t*-test was used for data analysis: **, 0.001 < p < 0.01; ns, no significant difference.

In every *mec-4(e1611)* mutant embryo, both the PLML and PLMR neurons undergo necrosis [19]. In worms that exhibit normal clearance of dying cells, these necrotic neurons are rapidly engulfed and degraded; as a consequence, some of them disappear before the embryo hatches [19,49]. To allow necrotic cells to persist during the larval stages, we included a *ced-1(e1735)* null mutation, which perturbs necrotic cell clearance [19], in our strains of interest. In each *ced-1(e1735); mec-4(e1611)* double mutant L1 larva hatched within 1 hr, on average 1.7 instead of 2 necrotic PLMs were observed (**Fig 3C**), likely due to the residual cell corpse clearance activity that is still present in *ced-1* mutants [50,51]. We found that low doses of dantrolene (e.g. 1.0 and 2.0 μM) did not significantly affect the number of PLM neurons that undergo necrosis (**Fig 3C**), yet resulted in a ∼40% reduction of PS level on the surface of necrotic PLMs (**Fig 3 A and B(d-i)**). This suggests that a modest level of inhibition of the release of Ca^2+^ from the ER affects the exposure of PS on necrotic touch neurons without significantly inhibiting necrosis. At higher doses (e.g. 3.0 and 4.0 μM), dantrolene resulted in a partial suppression of necrosis (∼32% reduction of the number of necrotic cells) (**Fig 3C**)) and a more severe reduction of the PS intensity (**Fig 3 A-B**). Collectively, these results suggest that while the Ca^2+^ released from the ER helps to induce both necrosis and PS exposure, PS exposure requires a higher level of cytoplasmic Ca^2+^ than cell swelling does. Moreover, they suggest that cytoplasmic Ca^2+^ might directly induce PS exposure independently of other features of necrosis.

To examine whether the dantrolene treatment indeed reduces the concentration of cytoplasmic Ca^2+^ in the PLM neurons in *mec-4(e1611)* mutant embryos, we monitored the GCaMP5G reporter over time. We first measured the Ca^2+^ levels in living PLMs whose necrosis was suppressed by 4 μM dantrolene and compared them to their untreated counterparts, and found that dantrolene treatment greatly reduced the Ca^2+^ levels in the cytoplasm of PLM neurons throughout embryonic development **(Fig 4)**. Remarkably, dantrolene treatment disrupted the transient cytoplasmic Ca^2+^ peak characteristic of necrosis **(Fig 4 a(o-bb))**. Secondly, we compared the cytoplasmic Ca^2+^ levels in necrotic PLM neurons in worms either through 3 μM dantrolene treatment or not treated with dantrolene. After 3 μM dantrolene treatment, the GCaMP5G signal intensity in necrotic PLM neurons display peaks that are lower and more ephemeral than their untreated counterparts **(Figs 4B and S1(C-D))**. Together, the heavily suppressed cytoplasmic Ca^2+^ signal patterns demonstrate that dantrolene treatments indeed cause the reduction of cytoplasmic Ca^2+.^ in the *mec-4(d)* background. These patterns correlate with the reduced PS level on the surface of necrotic PLM neurons and the partial suppression of necrosis, which is represented by cell swelling **(Fig 3)**.

**Figure 4.**
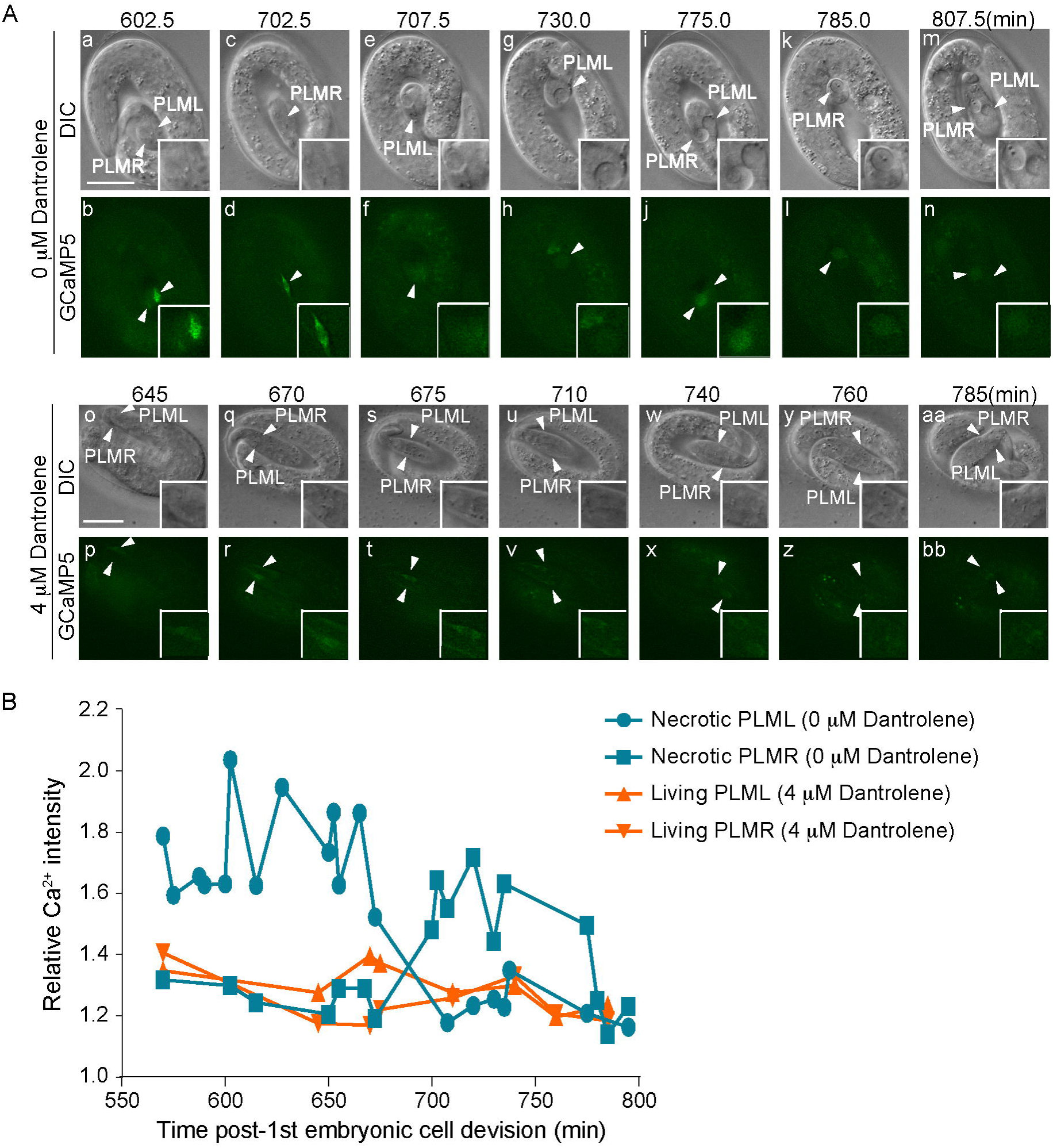
Dantrolene reduces the level of Ca^2+^ in the cytoplasm of touch neurons. Results of time-lapse recording experiments measuring the intensity of cytoplasmic Ca^2+^ and cell swelling of PLM neurons over time. The strain is homozygous for *mec-4(e1611)* allele and carries a transgenic array expressing P_*mec-7*_ *GCaMP5G::gfp*. (A) Time-lapse recording images of two necrotic PLM neurons in one untreated embryo (a-n) and two living PLM neurons in one embryo from the 4*µ*M dantrolene treated adult (o-bb). (a-n) Because PLML and PLMR do not appear in the same focal planes, at every time point only one of the two (identity labeled) is visible. (o-bb) No PLM swelling was observed in this embryo. Scale bars are 10*µ*M. (B) The relative fluorescence intensities of cytoplasmic GCaMP5 in the PLM neurons shown in (A) are plotted over time.

### Impairing the establishment of the Ca^2+^ pool in the ER inhibits PS exposure on necrotic neurons

To further dertermine the role of the ER Ca^2+^ pool in inducing PS exposure, we analyzed PS exposure on necrotic neurons in *crt-1* (*lf*) mutants in which the establishment of Ca^2+^ pool in the ER is defective. *crt-1* encodes the only *C. elegans* homolog of mammalian calreticulin, an ER-localized calcium chaperon protein that binds free Ca^2+^ and enables the accumulation of Ca^2+^ in the ER [52]. Calreticulin-deficient cells are defective in the storage of Ca^2+^ in ER [53]. In *C. elegans, lf* mutations in *crt-1* suppress the necrosis of neurons induced by dominant mutations in multiple DEG/ENaC genes, including *deg-1* [35]. In *deg-1(u38ts)*; *crt-1(lf)* double mutants and *deg-1(u38ts)* single mutant worms, we examined PS exposure on necrotic neurons using the PS reporter P_*ced-1*_*mfg-e8::mKate2* (**Materials and Methods**). In *deg-1(u38ts)* mutants, a small number of sensory and interneurons undergo necrosis at 25°C, the restrictive temperature [44], and expose PS on their surfaces (**Fig 5 A and B(b)**). Two *lf* alleles of *crt-1, bz29* and *bz50*, partially suppress necrosis and reduce the number of necrotic cells in *deg-1(u38ts)* mutants (**Fig 5C** and [35]). In both *crt-1* alleles, we found that the levels of PS signal on the existing necrotic cells were reduced to <10% of that in the *crt-1(+)* animals (**Fig 5 A and B(c-f)**). These results strongly support our model that a normal ER pool of Ca^2+^ and the release of Ca^2+^ into the cytoplasm are essential for proper PS exposure on necrotic cells.

**Figure 5.**
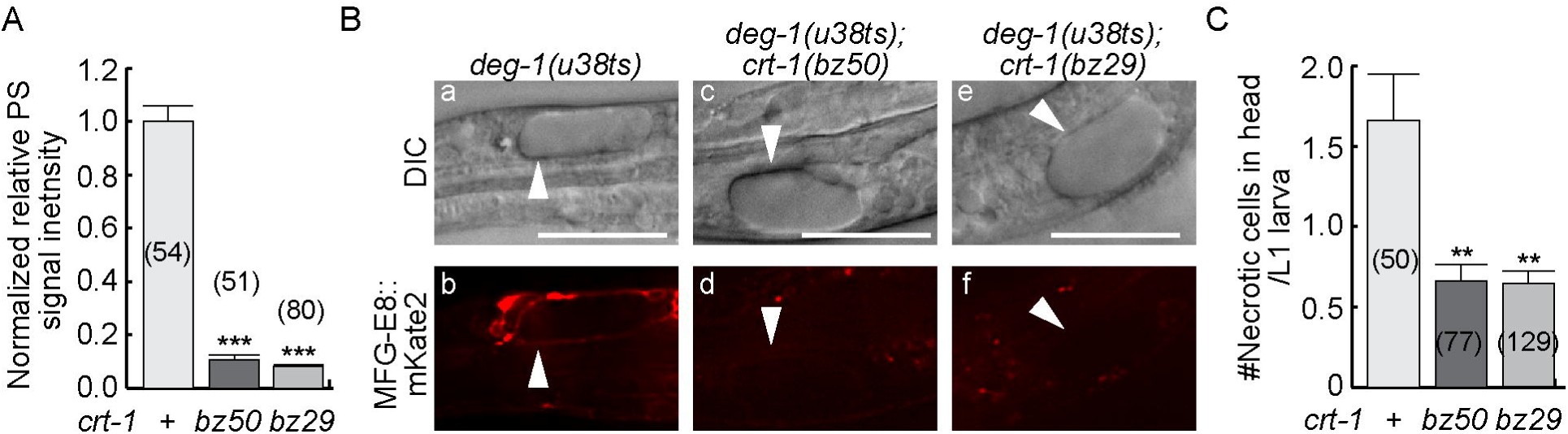
Mutations in *crt-1* strongly inhibit the exposure of PS on the surface of necrotic cells in *deg-1* dominant mutant animals. Newly hatched L1 larvae expressing the PS reporter P_*ced-1*_*mfg-e8::mKate2* are scored for the relative PS intensity (A-B) and the number of necrotic cells (C). All strains carry the *deg-1(u38ts)* mutation and were incubated at 25°C. (A) The normalized relative signal intensities of MFG-E8::mKate2 on the surfaces of necrotic cells in the heads are presented as the mean ratios relative to *crt-1(+)* worms. Bars represent the mean values of each sample. Error bars represent s.e.m.. Numbers in parentheses represent the number of animals analyzed. The Student *t*-test was used for statistical analysis. “***”, p<0.001. (B) DIC (a, c, e) and mKate2 fluorescence (b, d, f) images of necrotic cells (arrowheads) in the heads of newly hatched L1 larvae. Scale bars are 15*µ*m. (C) The mean numbers of necrotic cells in the head of each newly hatched larva are represented as bars. Error bars represent s.e.m.. Numbers in parentheses indicate the number of animals analyzed for each strain. The Student *t*-test was used for statistical analysis. “**”, <0.001<p<0.01, Student *t*-test.

### Ectopically increasing cytoplasmic Ca^2+^ induces necrosis and the exposure of PS

If the level of cytoplasmic Ca^2+^ determines whether a cell expose PS and undergo necrosis or not, presumably an ectopic increase of cytoplasmic Ca^2+^ level will induce cells to expose PS. To test this possibility, we treated wild-type worms with thapsigargin (**Materials and Methods**). Thapsigargin inhibits <u>S</u>arco-<u>E</u>ndoplasmic <u>R</u>eticulum <u>C</u>a^2+^ <u>A</u>TPase (SERCA), a Ca^2+^ re-uptake pump located on the ER and sarcoplasmic reticulum (SR) that pumps Ca^2+^ from the cytoplasm to the ER and SR [54,55]. Inhibiting SERCA activity increases the level of cytoplasmic Ca^2+^ [55]. We observed that, consistent with the previously report [35], thapsigargin treatment induced the onset of necrosis of cells that are otherwise destined to live in the head (**Fig 6 A-B**), whereas untreated worms exhibited nearly no necrotic cells (only one necrotic cell observed in a popuation of 83 larvae) (**Fig 6A**). Furthermore, thapsigargin-induced necrotic cells expose PS on their surfaces, with a higher concentration of thapsigargin correlating with stronger PS exposure (6 μg/ml vs 3 μg/ml thapsigargin) (**Fig 6 B-C**). These results thus support a model proposing that the PS exposure on cell surfaces is dependent on the concentration of cytoplasmic Ca^2+^.

**Figure 6.**
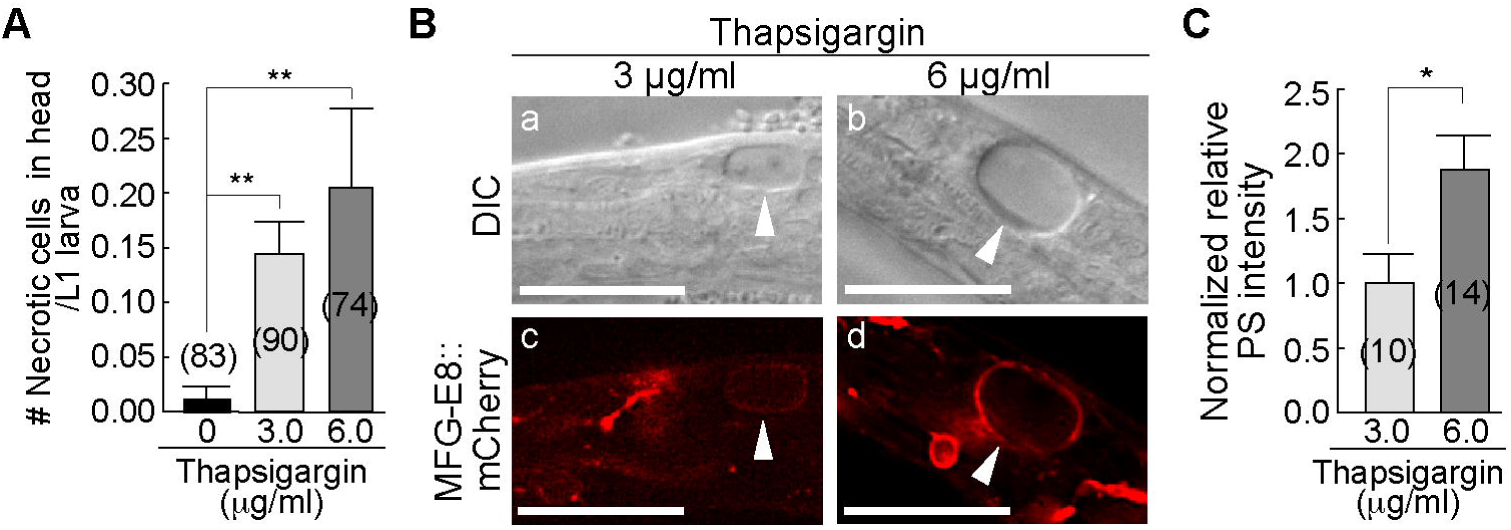
Thapsigargin induces live cells to undergo necrosis and expose PS on their surfaces. Adult worms carrying the integrated PS reporter P_*ced-1*_ *mfg-e8::mKate2* were placed on NGM plates with different doses of thapsigargin and incubated at 25°C; their progeny were allowed to hatch into L1 larvae and promptly scored (within 1 hr after hatching) for the number of necrotic corpses in the head (A) and the intensity of PS on the surface of necrotic cells (B and C). (A) The mean numbers of necrotic cells in each L1 larva are represented in the bar graph. Error bars represent s.e.m.. “**”, 0.001<p<0.01, Student *t*-test. (B) DIC (a, b) and MFG-E8::mCherry (c, d) fluorescent images of necrotic cells (white arrowheads) observed in the head. Scale bars are 15*µ*m. (C) The normalized relative MFG-E8::mCherry signal intensities on the surfaces of necrotic cells in the heads are presented as the mean ratios relative that measured from worms subject to 3 μM thapsigargin treatment. Error bars represent s.e.m.. Numbers in parentheses represent the number of animals analyzed for each strain. “*”, 0.01<p<0.05, Student *t*-test.

### Induction of PS exposure is independent of the induction of other necrosis features

Our observation that low concentrations of dantrolene are able to suppress PS exposure but not the necrotic cell swelling in the *mec-4(e1611)* mutant animals **(Fig 3)** suggests that the relationship between PS exposure and increased cytoplasmic Ca^2+^ might exist independently of other events occurring during necrosis. We reason that if PS exposure is a consequence of the execution of necrosis, blocking the intermediate steps of necrosis execution would inhibit PS exposure even when cytoplasmic Ca^2+^ is increased via ER-mediated Ca^2+^ release; otherwise, PS exposure will not be blocked but other necrotic cell events such as cell swelling will. We applied two different methods to suppress the intermediate necrosis events, and examined whether PS exposure is affected in each case.

We first explored the effect of an *unc-51(e369)* mutation on PS exposure. *unc-51* encodes a homolog of yeast ATG1 and mammalian ULK1 (<u>U</u>NC-51-like autophagy activating <u>k</u>inase), an important component of the autophagy pathway [56]. Autophagy genes actively participate in the lysosomal-dependent necrosis, which causes cell swelling and other cellular events [57,58]. Specifically, a loss-of-function mutation of *unc-51* was reported to partially block the execution of necrosis induced by a *mec-4(d)* allele [57,58]. We introduced a *unc-51(e369)* mutant allele into the *ced-1(e1735); mec-4(e1611)* mutant strain that carries the MFG-E8::mCherry and P_*mec-7*_*gfp* reporters. We specifically monitor the PLML and PLMR neurons in the tail of young L1 larvae. Consistent with previous reports [57,58], we observed a significant and sizable reduction (25%) in the number of necrotic PLM neurons in newly hatched *ced-1(e1735); unc-51(e369); mec-4(e1611)* L1 larvae comparing to that observed in *ced-1(e1735); mec-4(e1611)* L1 larvae (**Fig 7A**). Remarkably, PS is detected on 27.4% of the PLM neurons (labeled with P_*mec-7*_ GFP) that appear live and display a relative normal and non-swelling morphology under the DIC microscope in *ced-1(e1735); unc-51(e369); mec-4(e1611)* L1 larvae (**Fig 7 B-C)**, whereas in *ced-1(e1735)*; *unc-51(+); mec-4(+)* L1 larvae in which all PLM neurons are alive, 0% of living PLM neurons expose PS (**Fig 7 B-C** and [19]). **Fig 7B(a-c)** displays an example of an apparently living PLM neuron that exposes PS. This phenomenon reveals that PS exposure could be induced by the *mec-4(d)* mutation in a manner independent of necrotic cell swelling.

**Figure 7.**
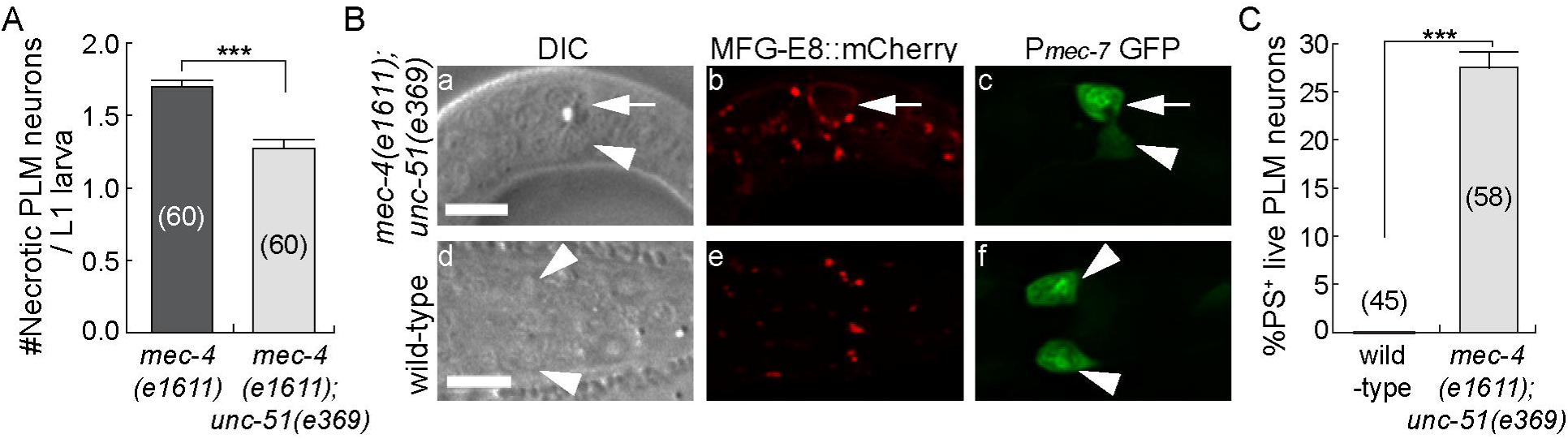
Impairing the execution of necrosis does not block PS exposure. (A) The mean number of PLM neurons (labeled with the P_*mec-7*_ GFP reporter) in *ced-1(e1735); mec-4(e1611)* and *ced-1(e1735); unc-51(e369); mec-4(e1611)* L1 larvae that displayed the necrotic swelling phenotype were presented in the graph. L1 larvae were scored within 1 hr of hatching. Bars represent the mean values of each sample. Error bars represent s.e.m.. The numbers in the parentheses represent the numbers of animals scored. “***”, p<0.001, Student *t*-test. (B) DIC, GFP, and mCherry images of four living PLM neurons, two in a *ced-1(e1735); unc-51(e369); mec-4(e1611)* L1 larva (a-c, white arrows and arrowheads) and two in a wild-type L1 larva (d-f, white arrowheads). PLM neurons are identified by the P_*mec-7*_GFP reporter (c, f). DIC images (a, d) show that none of the four PLM neurons display the necrotic swelling morphology. PS presentation (MFG-E8::mCherry) on the surface of one PLM neuron (b) is marked by a white arrow (b, e). White arrowheads in (a-f) mark living PLM neurons that do not expose PS. Scale bars are 5*µ*m. (C) A bar graph representing the percentage of live PLM neurons that expose PS on their surfaces among all living PLM neurons in early L1 larvae with the indicated genotypes. Error bars represent s.e.m.. The numbers in the parentheses represent the numbers of animals scored. “***”, p<0.001, Student *t*-test.

We next attempt to impair the execution of necrosis through disrupting lysosomal function. During excitotoxic necrosis, increased cytoplasmic concentration of Ca^2+^ is known to activate a cascade of events that lead to the breakage of lysosomes, the acidification of the entire cell, and the release of lysosomal hydrolytic enzymes into the cytoplasm [59,60]. Cell swelling is a primary consequence of these events [59,60]. To disrupt lysosomal function, we applied NH_4_Cl, a reagent that impairs lysosomal acidification and partially blocks the execution of necrosis of touch neurons in the *mec-4(d)* mutants [59], to a liquid worm culture (**Materials & Methods**). In the *ced-1(e1735)*; *mec-4(e1611)* mutant animals, 5mM NH_4_Cl causes a 40% reduction in the necrosis of the PLM neurons in L1 larvae (**Fig S2A**), consistent with a previous report [59]. In some of those live (non-swelling) PLM neurons, we observed PS exposure on their cell surfaces (**Fig S2B(d-f)**, white arrows). Together, the results obtained from the *unc-51* mutants and the NH_4_Cl experiment indicate that the exposure of PS is not dependent on the full execution of necrosis.

### Impairing the release of Ca^2+^ into the cytoplasm does not affect PS exposure on the surfaces of apoptotic cells

During apoptosis in *C. elegans*, whether cytoplasmic Ca^2+^ contributes to the PS exposure on apoptotic cells was not known. To address this question, we examined whether a *crt-1(bz29)* null mutation, which disrupts the Ca^2+^ pool in the ER, would affect the PS exposure on apoptotic cells. In *ced-1(e1735)* null mutants, the clearance of apoptotic cells is largely inhibited and many apoptotic cells persist in the head of newly hatched L1 larvae (**Fig 8 A-B**). Loss of *ced-1* function does not affect PS dynamics on the surfaces of dying cells [19,20], and thus all of these apoptotic cells expose PS on their cell surfaces (**Fig 8 B-C**). In thw *ced-1(e1735); crt-1(bz29)* double mutant L1 larvae, the *crt-1(bz29)* null mutation does not affect either the number of apoptotic cells or the level of PS exposure on apoptotic cells **(Fig 8 A-C)**, in stark contrast to the effect of *crt-1(bz29)* on necrotic cells (**Fig 5**).

**Figure 8.**
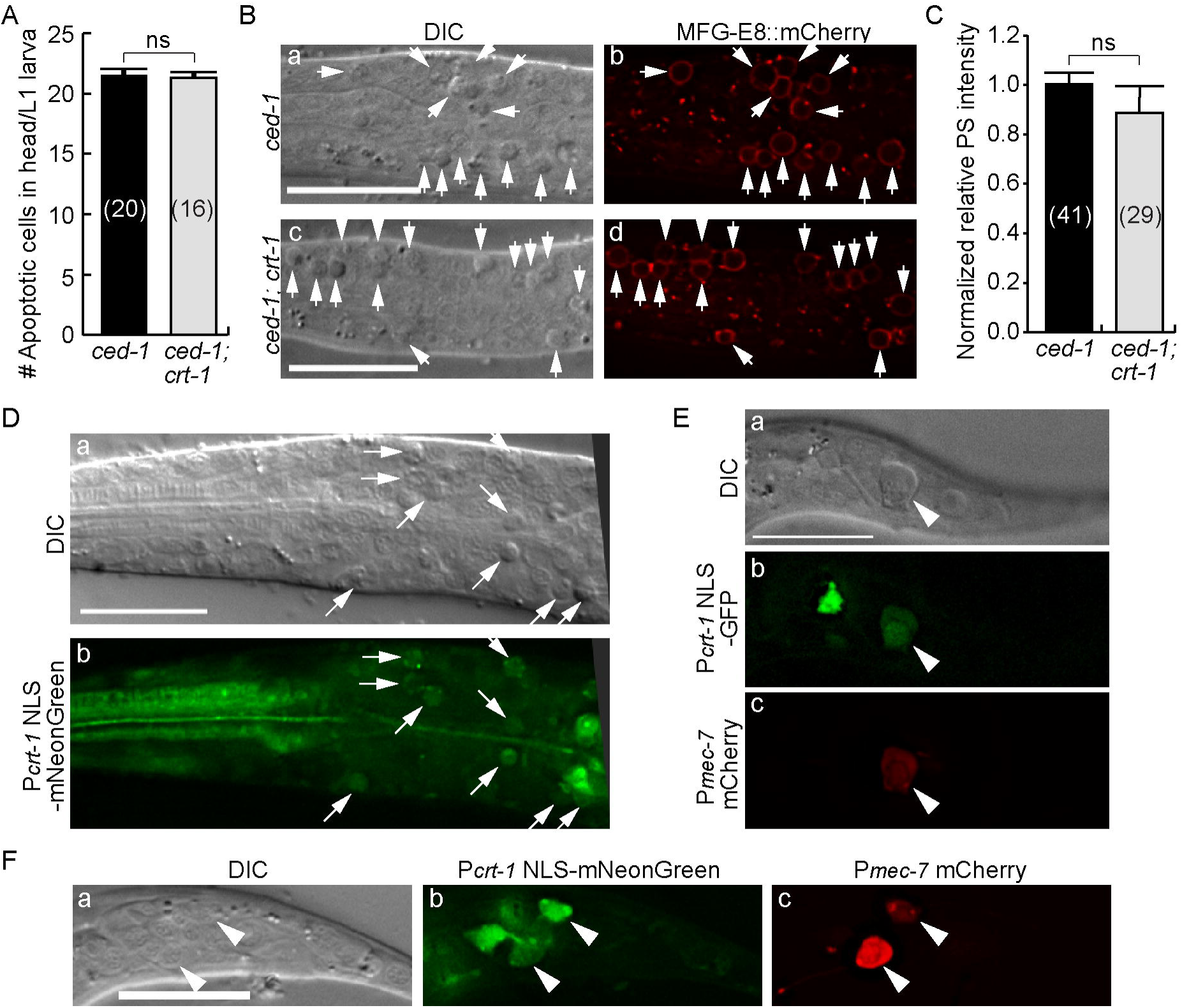
A null mutation in *crt-1* does not affect the exposure of PS on the surface of apoptotic cells. (A) In *ced-1(e1735)* and *ced-1(e1735); crt-1(bz29)* mutant strains, the number of apoptotic cells in the head of L1 larvae hatched within 1 hr were scored. Bars represent the mean values of each sample. Error bars represent s.e.m.. Numbers in parentheses represent the the numbers of L1 larvae scored. ns, statistically not significant (p>0.05, Student *t*-test). (B) DIC and fluorescence images of the heads of two larvae displaying numerous apoptotic cells (a and c, arrows). The PS signal exposed on their surfaces are detected by MFG-E8::mCherry (b and d, arrows). Scale bars are 10 *µ*m. (C) Apopotic cells in the heads of larvae were scored for the normalized MFG-E8::mCherry signal intensity on their cell surfaces. The signal intensity was compared between two different mutant strains. Bars represent the mean values of each sample. Error bars represent s.e.m.. Numbers in parentheses are the the numbers of apoptotic cells analyzed. ns, statistically not significant (p>=0.05, Student *t*-test). (D-F) The *crt-1* promoter is expressed in apoptotic cells (D), a necrotic touch neurons (E), and two living touch neurons (F). Transgenes were expressed in *ced-1(e1735); mec-4(e1611)* double (D and E) and *ced-1(e1735)* single (F) mutant strains. Scale bars are 15 *µ*m. (D) DIC (a) and fluorescence (b) images of the head of an L2/L3-stage larva expressing P_*crt-1*_NLS-mNeonGreen. Arrows mark apoptotic cells. (E) DIC (a) and fluorescence (b and c) images of the tail of an L1-stage larva co-expressing P_*crt-1*_NLS-GFP (b) and the touch neuron marker P_*mec-7*_mCherry (c). Arrowheads mark one necrotic PLM neurons. (F) DIC (a) and fluorescence (b and c) images of the tail of an L1-stage larva co-expressing P_*crt-1*_NLS-GFP (b) and P_*mec-7*_mCherry (c). Arrowheads mark living PLM neurons.

*crt-1* is broadly expressed in *C. elegans* [35,61]. We next determined whether *crt-1* is expressed in cells destined to undergo apoptosis and/or necrosis. We constructed P_*crt-1*_*NLS-gfp* and P_*crt-1*_*NLS-mNeonGreen* reporters, in which the GFP or mNeonGreen reporters are tagged with a nuclear localization signal (NLS) and expressed under the direct control of the *crt-1* promoter (P_*crt-1*_) [61] (**Materials & Methods**). The *crt-1* open reading frame is not present in these two reporters. The NLS sequence facilitates the enrichment of GFP or mNeonGreen in the nucleus. In the *ced-1(e1735); mec-4(e1611)* double mutant animals that express the P_*crt-*1_ *NLS-gfp* or P_*crt-*1_ *NLS-mNeonGreen* constructs, mNeonGreen and GFP signals were observed broadly, including in many apoptotic cells retained in the head (**Fig 8D**) as well as the necrotic touch neurons retained in the tail of L1 larvae due to the *ced-1* mutation (**Fig 8E**). In addition, in *ced-1(e1735)* single mutant larvae, P_*crt-1*_ NLS-mNeonGreen expression was observed in live PLM neurons in the tail (**Fig 8F**). These results indicate that *crt-1* is expressed in cells destined to die of apoptosis and necrosis. Therefore, Collectively, our observations support the hypothesis that the ER Ca^2+^ pool does not regulate either the initiation of apoptosis or the exposure of PS on apoptotic cells, despite that *crt-1* is expressed and presumably functional in cells distined to undergo apoptosis. Our findings thus demonstrate that necrotic and apoptotic cells employ different molecular mechanisms to trigger PS exposure.

## Discussion

Previously, the presentation of PS, the “eat me” signal, on the surfaces of necrotic cells was thought to be a result of the rupture of the plasma membrane and the passive exposure of the inner leaflet to the outside. We found that necrotic cells actively expose PS on their outer surfaces in order to attract engulfing cells [19]. Our continuing investigation reported here reveals a previously unknown signaling mechanism that triggers the exposure of PS on the surface of necrotic cells. Despite its important roles in many cellular events, Ca^2+^ was previously unknown to act as a trigger for the clearance of necrotic cells. We developed a time-lapse recording approach and monitored the dynamic change of the cytoplasmic Ca^2+^ concentration both prior to and throughout the necrosis of touch neurons in developing *C. elegans* embryos. The Ca^2+^ signal in touch neurons has previously been measured in adults [62] but has not been monitored in real time or during embryonic development. Our newly-developed real time recording protocol has revealed a rapid and transient increase of the cytoplasmic Ca^2+^ in the touch neurons prior to cell swelling, a feature of necrotic cells, and the presentation of PS on necrotic cell surface. Further quantitative analysis demonstrated a close correlation between the levels of cytoplasmic Ca^2+^ and PS exposure. We further discovered that the ER Ca^2+^ pool was necessary for the the robust increase in cytoplasmic Ca^2+^ and the consequential PS exposure during the excitotoxic necrosis of neurons. Based on these findings and on the known mechanisms that regulate the intracellular Ca^2+^, we propose a model describing an ER-assisted, Ca^2+^-triggered PS exposure-mechanism occuring in neurons undergoing necrosis (**Fig 9**).

**Figure 9.**
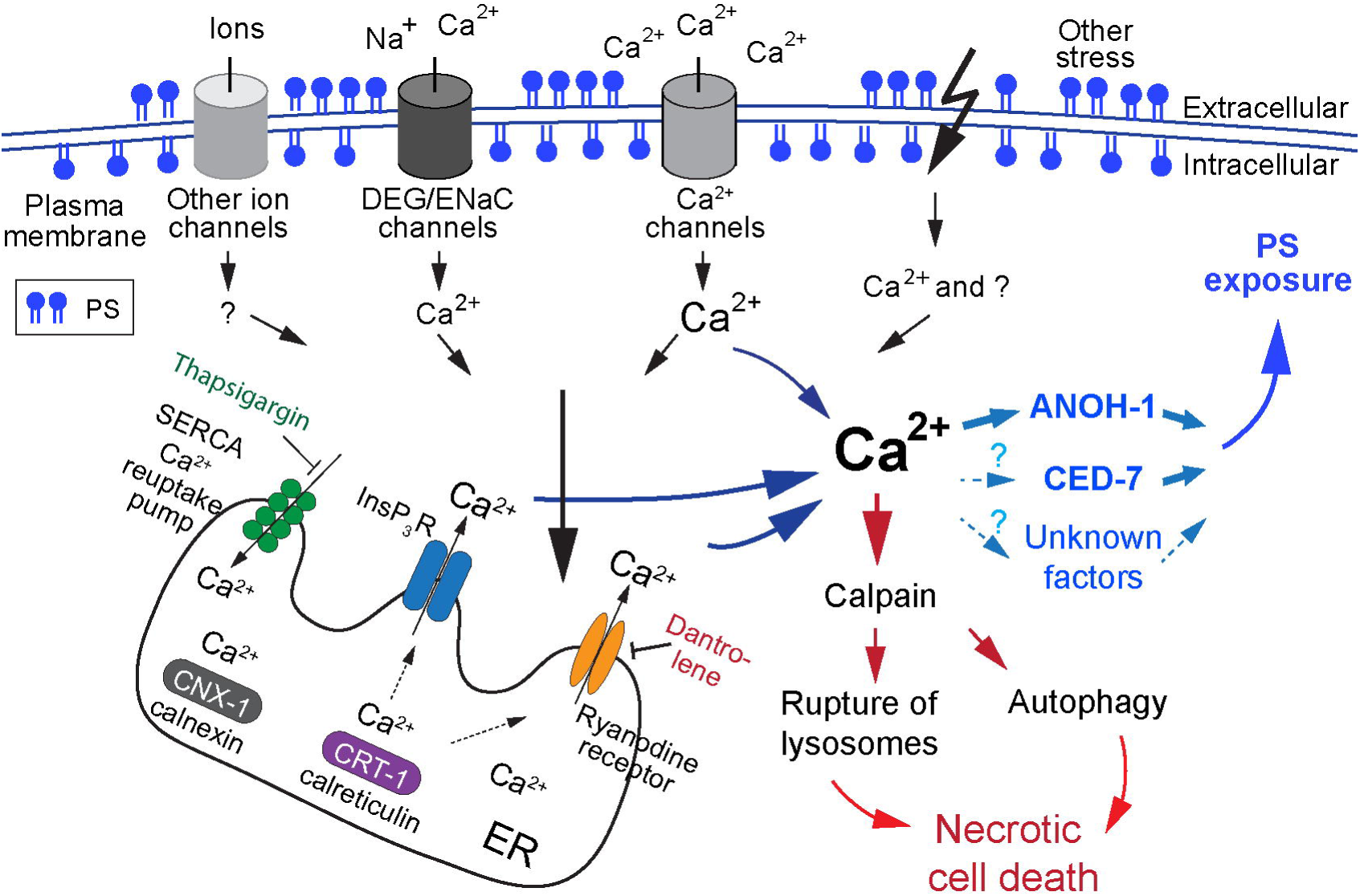
A model illustrating how the ER-assisted increase of cytoplasmic calcium ion induces the exposure of PS, the “eat me” signal, on the surface of necrotic cells. See Discussion for a more detailed explanation of the models. The cylinders on the plasma membrane represent various types of ion channels, which allow Ca^2+^ influx due to the mutations in certain subunits. The establishment of the calcium pool in the ER requires the calcium chaperons calreticulin (CRT-1) and calnexin (CNX-1). The release of Ca^2+^ from the ER requires the ryanodine receptor, whose activity is inhibited by dantrolene, and the InsP_3_ receptor (InsP_3_R). SERCA is an ER-surface Ca^2+^ reuptake pump whose function is inhibited by thapsigargin. Dominant mutations of channel subunits allow a small amount of Ca^2+^ to enter neurons. This, in turn, triggers a “Ca^2+^- induced” Ca^2+^ release from the ER, results in a further increase of cytoplasmic Ca^2+^ concentration. The resulting surge in the level of the cytoplasmic Ca^2+^ activates multiple downstream targets to induce parallel necrosis events. The Ca^2+^-dependent protease calpain is known to promote lysosomal rupture and autophagy. We propose that ANOH-1 is a prime candidate for a Ca^2+^-target that facilitates PS exposure. In addition, cytoplasmic Ca^2+^ might also activate CED-7 or other unknown PS-externalization enzymes (cyan question marks). Besides Ca^2+^, there might be other upstream signaling molecules (black question marks) that induce PS exposure. On the other hand, a strong influx of Ca^2+^ through hyperactive Ca^2+^ channels might induce PS exposure and necrosis in an ER-independent manner.

### An ER-assisted, Ca^2+^-triggered mechanism that promotes the exposure of PS on the surface of necrotic neurons

The DEG/ENaC sodium channel - in which MEC-4 is a subunit of - plays an essential role in the mechanosensory function of *C. elegans* touch neurons [13]. Biochemical studies have revealed that the MEC-4(d) mutant protein alters the property of this sodium channel, making it permeable to Ca^2+^ [36]. This altered permeability was detected when overexpressed in Xenopus oocytes or in *C. elegans* touch neurons, although only a small current of Ca^2+^ was detected [36]. In *mec-4(d)* mutants, multiple lines of evidence indicate that an increased level of cytoplasmic Ca^2+^ provided by the calcium-induced ER Ca^2+^ release mechanism triggers necrosis [35,38].

We have found that the exposure of PS is a common feature observed on multiple types of necrotic neurons (**Fig 1**). Our time-lapse observation of Ca^2+^ signal intensity in *mec-4(d)* mutants has provided direct evidence of a transient increase in the cytoplasmic Ca^2+^ level prior to necrotic cell swelling and PS exposure. We further provided multiple lines of evidence to indicate that, similar to cell swelling, the exposure of PS on the surface of necrotic neurons in *mec-4(d)* and *deg-1(d)* mutants is also regulated by the ER Ca^2+^ release: the chemical treatment that inhibit the release of Ca^2+^ from the ER and mutations that impair the establishment of the Ca^2+^ pool in the ER both attenuate PS exposure; on the contrary, the treatment that inhibits re-uptake of Ca^2+^ into ER not only induces necrosis of cells in otherwise wild-type animals, but also triggers the exposure of PS on the surfaces of these necrotic cells. These results alone may suggest that the exposure of PS exposure is merely a consequence of necrosis, but our further experiments demonstrate that this is not the case.

Firstly, while low doses of dantrolene reduce the intensity of PS on the necrotic cell surfaces without suppressing the frequency of cell swelling, higher doses both partially suppress necrosis and further reduce PS intensity on the surfaces of the remaining necrotic cells **(Fig 3)**. This phenomenon suggests that the threshold of cytoplasmic Ca^2+^ required for initiating cell swelling is lower than that needed for inducing the full blown presentation of PS, suggesting that cell swelling and PS exposure might be regulated by different calcium effectors. Secondly, we directly examined whether PS exposure is a downstream consequence of necrosis by blocking *mec-4(d)*- induced necrosis, through either inhibiting autophagy or impairing proper lysosomal function. Both of these approaches were able to suppress cell swelling, generating normal-looking touch neurons. Remarkably, even in the absence of cell swelling, PS was frequently detected on the plasma membrane of these normal-looking neurons. These observations indicate that the Ca^2+^-induced PS-exposure can be blocked independently of cell swelling (and possibly other features of necrosis), strongly suggest that the cytoplasmic Ca^2+^ targets multiple effectors in parallel, one of which leads to PS exposure.

Although we have found a correlation between the increase of cytoplasmic Ca^2+^ levels and PS exposure, the mechanisms behind this phenomenon are not yet known. One potential candidate is TMEM16F, the mammalian homolog of ANOH-1. TMEM16F is both a Ca^2+^ channel and a Ca^2+^-dependent phospholipid scramblase [24,25]. Although its biochemical activity has not been determined, *C. elegans* ANOH-1 might act as a phospholipid scramblase whose activity is dependent on its association with Ca^2+^. In this scenario, the ER-dependent increase of cytoplasmic Ca^2+^ level induced by the *mec-4(d)* mutation might directly activate ANOH-1, which in turn promotes PS exposure (**Fig 9**). Conversely, CED-7, the other protein important for PS exposure, is not known to bind Ca^2+^. It remains to be investigated whether CED-7 is also regulated by Ca^2+^ or by a different upstream signaling molecule. Given that CED-7 is important for PS exposure on the surface of both apoptotic and necrotic cells [19,20], it is possible that CED-7 can be activated by multiple upstream signals.

As for how ER Ca^2+^ mediates cell swelling, the Ca^2+^-activated protease calpain is a known target of necrosis signals, whose stimulated activity is associated with lysosomal rupture and the activation of autophagy, two major events that lead to many cell morphological changes and degradation of cellular proteins during necrosis [63-66]. We propose that during the initiation of necrosis, the elevated cytoplasmic Ca^2+^ concentration activates, in parallel, a number of downstream targets that include the PS-exposure enzymes, calpain, and possibly other proteases and results in cellular changes of multiple aspects, including PS exposure and cell swelling (**Fig 9**).

### The possibility of a Ca^2+^-dependent, ER-independent trigger of PS exposure

The MEC-4(d) mutations that induce necrosis only allows a very modest level of Ca^2+^ permeability through the touch neuron-specific DEG/ENaC sodium channels [36]. This might be why the release of Ca^2+^ from the ER is necessary to further increase the cytoplasmic Ca^2+^ level over the threshold required for necrosis and PS exposure. Similarly, the necrosis and PS exposure triggered by a dominant mutation in *deg-1*, which encodes another DEG/ENaC family sodium channel subunit, also depends on the contribution of the ER. However, it would stand to reason that if the Ca^2+^ influx from the extracellular space is strong enough, the ER-dependent Ca^2+^-release could be dispensible for PS exposure. Among the insults that are known to trigger the necrosis of neurons, a dominant mutation in *deg-3*, which encodes a subunit of a ligand-gated calcium channel belonging to the nicotinic acetylcholine receptor family [43], induces neuronal necrosis in a ER-independent manner [35]. The *deg-3(u662)* dominant mutation results in a calcium channel with increased conductivity [43,67]. It is proposed that a large amount of Ca^2+^ might consequentially enter neurons from the extracellular space, large enough to trigger necrosis without the aid of additional Ca^2+^ released from the ER [35]. We propose that likewise, when the influx of Ca^2+^ from extracellular space is large enough, the Ca^2+^- triggered PS exposure might be independent of the ER Ca^2+^ release.

### Does intracellular Ca^2+^ facilitate PS exposure on the surface of apoptotic cells?

Ca^2+^ is a well-known trigger for PS exposure on the surface of platelets for the initiation of blood co-aggulation [68]. Previously, extracellular Ca^2+^ was reported to stimulate the exposure of PS on the surface of a T lymphocyte hybridoma cell line undergoing apoptosis [69]. A calcium-dependent PS exposure mechanism was suggested to be a general mechanism utilized by apoptotic cells of different identities [70]; however, there are lines of evidence that do not agree with this proposal [71,72]. Nonethless, it was unknown whether Ca^2+^ is involved in the developmentally programmed apoptosis and/or the subsequent exposure of the “eat me” signal in *C. elegans*. We determined that neither the number of apoptosis events nor the level of PS exposure is compromised during embryogenesis in *crt-1* mutants, which exhibit defects in establish the ER Ca^2+^ pool. These results indicate that the fluctuation of the cytoplasmic Ca^2+^ level does not trigger either apoptosis or PS exposure in developing *C. elegans* embryos. This is consistent with our previous finding that the proposed Ca^2+^-dependent phospholipid scramblase ANOH-1 is not involved in the PS exposure on apoptotic cells [19]. Together, our findings demonstrate that apoptotic and necrotic cells regulate the exposure of “eat me” signals and their subsequent clearance through distinct mechanisms; apoptotic cells utilize Ca^2+^-independent mechanisms, while necrotic cells utilize Ca^2+^-dependent ones.

### The Ca^2+^-dependent regulatory mechanism for PS exposure is likely conserved throughout evolution

The regulatory mechanism we have discovered in *C. elegans* might be a mechanism that is also employed in other organisms including mammals, in particular, since the homolog of mammalian Ca^2+^-dependent scramblase TMEM16F not only exists in *C. elegans*, but also in other organisms as well. Whether TMEM16F plays a similar role as *C. elegans* ANOH-1 in facilitating the clearance of necrotic cells in mammals remains to be tested, although it is clear that TMEM16F is not involved in the PS exposure on apoptotic cells [73]. Insults that increase the cytoplasmic Ca^2+^ trigger the necrosis of neurons, glial cells, cardiomyocytes, cancer cells, and other cells [6,34,74-76], and it is possible that these necrotic cells all utilize the Ca^2+^-dependent mechanism to activate the PS-exposure activity. We predict that the Ca^2+^-triggered PS exposure might be an evolutionarily conserved mechanism that facilitates the clearance of different types of cells that undergo necrosis in a wild variety of organisms. As calcium overload and the consequential excitotoxic necrosis play critical roles in the pathology of many diseases including neurodegenerative disorders, stroke, heart failure, and cancer [64,74-76], investigating the clearance of necrotic cells using *C. elegans* as a model organism will shed light on the pathology and therapeutics of these human diseases. For example, there are pre-clinical therapeutic attempts to use chemical inhibitors of the Ca^2+^-dependent protease calpain to hamper the necrosis occurring during pathological conditions such as brain ischemia [77]. What we have found suggests that this approach might no be able to efficiently inhibit the exposure of PS on cells whose necrosis is blocked and thus dose not necessarily prevent these cells from being engulfed.

## Materials and Methods

### Mutations, plasmids, strains, and transgenic arrays

*C. elegans* was grown at 20°C as previously described [78] unless indicated otherwise. The N2 Bristol strain was used as the wild-type strain. Mutations are described in [79] and in the Wormbase (http://www.wormbase.org) unless noted otherwise: LGI, *ced-1(e1735), trp-4(ot337)*; LGII, *enIs74*[P_*dyn-1*_*mfg-e8::mCherry*, P_*mec-7*_ *gfp*, and *punc-76(+)*] (this study); LGIII, *crt-1* (*bz29 and bz50*); LGIV, *unc-8(n491sd);* LGV, *unc-76(e911), deg-3(u662), unc-51(e369), enIs92*[P_*mec-7*_*GCaMP5G*, P_*dyn-1*_*mfg-e8::mCherry*, and *punc-76(+)*] (this study); LGX, *deg-1(u38ts), mec-4(e1611)*.

Plasmids P_*dyn-1*_*mfg-e8::mCherry* and P_*mec-7*_*gfp* (pPD117.01, a gift from Andrew Fire) have been reported in [19]. P_*ced-1*_*mfg-e8::mKate2* was generated by first replacing the *dyn-1* promoter (P_*dyn-1*_) [80] in P_*dyn-1*_*mfg-e8::mCherry* with P_*ced-1*_ [23] and then replacing mCherry cDNA in P_*ced-1*_*mfg-e8::*mCherry with mKate2 cDNA [81]. P_mec*-7*_*GCaMP5G* was generated by cloning the GCaMP5G cassette obtained from pCMV-GCaMP5G [47] to the BamH1 and EcoR1 sites of pPD117.01. Plasmid P_*crt-1*_*NLS-gfp* was generated by amplifying the genomic DNA that covers the first 6 amino acids including the start codon and a 1.5kb upstream region representing the *crt-1* promoter (P_*crt-1*_) [82] and cloning this fragment into the Sph1 and BamH1 sites of pPD95.69 (a gift from Andrew Fire), in frame with the nuclear localization signal (NLS) tagged GFP coding sequence, generating a NLS-GFP reporter expressed under the control of P_*crt-1*_. Plasmid P_*crt-1*_*NLS-mNeonGreen* was constructed by replacing the coding sequence for GFP with that for mNeonGreen [83].

Extrachromosomal arrays were generated by microinjection [84] of plasmids with coinjection marker p*unc-76(*+) [85] into strains carrying the *unc-76(e911)* mutation. Transgenic animals were isolated as non-Unc animals. *enIs74* and *enIs92* are integrated transgenic arrays generated from the corresponding transgenic arrays in this study. *trp-4(ot337), crt-1*(*bz29), crt-1(bz50), unc-8(n491sd)*, and *deg-1(u38)* were provided by the *C. elegans* Genetic Center (CGC).

### Chemical treatments of *C. elegans*

#### Dantrolene and thapsigargin treatments

Worms were treated with various dose of dantrolene (Tocris Bioscience, Inc.) as described previously [35]. Dantrolene was dissolved in DMSO and was both spread on unseeded NGM plates and added to the suspension of OP50 (the *E. coli* strain seeded onto the NGM plates). One day after seeding the NGM plates with the drugged OP50, L4 hermaphrodites were placed onto the dantrolene plates and raised at room temperature (20-22°C). Twenty-four to 36 hrs later, L1 larvae (F1 progeny) hatched within 1 hr were examined by both DIC and fluorescence imaging. Dantrolene is likely affecting embryos through entering the germline of adult hermaphrodites and subsequently entering the embryos generated in the germline. The 0 μM dantrolene control sample represent worms from NGM plates and seeded OP50 that were treated with DMSO, the solvant for dantrolene, and no dantrolene. The thapsigargin (Sigma-Aldrich, Inc.) treatment protocol is similar to that of dantrolene treatment, except that the worms were raised at 25°C instead of room temperature (20-22°C).

#### NH_4_Cl treatment

As described previously [35], 1M NH_4_Cl (Sigma-Aldrich, Inc.) solution in water was added to 10 mL S medium containing 50 young adult hermaphrodites and appropriate amount of concentrated bacteria OP50 (food for worms) to reach a final concentration of 5 mM. This suspension was grown at room temperature and under constant shaking. Fifteen hours later, young L1 larvae were collected and analyzed by DIC and fluorescence microscopy. Note that because the progeny of the young adults were grown in the liquid culture and could not be distinctly timed, the L1 larvae that we collected and analyzed are between 1-3 hrs after hatching. As a consequence, the mean number of necrotic PLMs observed in the tail is slightly lower than that of the L1 larvae scored within 1 hr of hatching due to the extra time after hatching.

### DIC microscopy and scoring the number of necrotic cells and apoptotic cells

DIC microscopy was performed using an Axioplan 2 compound microscope (Carl Zeiss) equipped with Nomarski DIC imaging apparatus, a digital camera (AxioCam MRm; Carl Zeiss), and imaging software (AxioVision; Carl Zeiss), or with an Olympus IX70-Applied precision DeltaVision microscope equipped with a DIC imaging apparatus, a Photometrics Coolsnap 2 digital camera, and the SoftWoRx imaging software (GE Healthcare, Inc.). In *ced-1(e1735); mec-4(e1611)* mutant and *ced-1(e1735); mec-4(+)* control backgrounds, the presence of two necrotic PLM neurons (PLML and PLMR) in the tail was scored by their swelling morphology in L1 larvae as previously described [86]; these larvae were collected within 1 hr after hatching, or within 1-3hrs after hatching, depending on what is annotated in the figure legends. Between 30 - 60 L1 larvae were scored for each sample. These larvae were grouped as 10 animals per group. The mean values of the number of necrotic PLM neurons of each group were calculated, and the mean values and standard errors of the means of all groups were presented in bar graphs. To examine *deg-1(u38ts)*-induced necrosis, adults were grown at 25° C for one day, at which point we collected the embryos that they laid. These embryos were incubated at 25°C and allowed to hatch, and we examined the L1 larvae within 1hr after they hatched. Between 50 - 129 larvae were scored for the number of necrotic cells in the head region. These larvae were grouped as 10, 15, and 25 animals per group for the *deg-1(u38ts), deg-1(u38ts); crt-1(bz50)*, and *deg-1(u38ts); crt-1(bz29)* strains, respectively. The mean values of the number of necrotic cells of each group were calculated, and the mean values and standard errors of the means of all groups were presented in bar graphs.

The number of apoptotic cells was scored in the head of newly hatched L1 larvae as described in [86].

### Fluorescence microscopy, time-lapse recording, and quantification of image intensity

An Olympus IX70-Applied Precision DeltaVision microscope equipped with a DIC imaging apparatus and a Photometrics Coolsnap 2 digital camera was used to acquire serial Z-stacks of fluorescence and DIC images, whereas the SoftWoRx software was utilized for image deconvolution and processing as described in [19,86]. To quantify the intensity of PS signal on the surface of a necrotic PLM neuron, the cell surface area containing the MFG-E8::mCherry signal was outlined by two closed polygons and the PS signal intensity (PS_PLM_) of the “donut shape” area between the two polygons was calculated as the signal value of the bigger polygon subtracted by that of the smaller polygon. Likewise, the background mCherry signal (PS_Background_) of a “donut shape” area in equivalent size was obtained by copying pasteing the two aforementioned polygons to a region neighboring the necrotic neuron. The relative PS signal intensity (PS_R_) = PS_PLM_ / PS_background_. The PS intensity on the surfaces of other necrotic neurons, living PLM neurons, and apoptotic cells are also measured using the above method.

To quantify the Ca^2+^ intensity in the cytoplasm of live or necrotic PLM neurons, the cell body of a PLM neuron was outlined by a polygon and the GCaMP5G signal value within the polygon was obtained and referred to as Ca_x_. Ca_Background_ of an area in equivalent size in a neighboring region was similarly obtained. The relative Ca^2+^ intensity of a particular cell was defined as Ca_R_ = Ca_x_ / Ca_Background_.

To monitor the dynamics of Ca^2+^ release into the cytoplasm of PLM neurons, cell swelling during necrosis, and PS presentation during embryogenesis, we modified a previously established protocol [19]. We mounted embryos on an agar pad on a glass slide with M9 buffer. The recording period covered 560 min (100 min after worms reached the 2-fold stage (460min)) to 900 min post-1^st^ embryonic cleavage. Serial Z stacks were captured in 10-15 sections at 0.5*µ*m per section. Recording interval was 2.5 min from 560 - 680 min post-1^st^ embryonic cleavage and 5 min from 680 - 900 min post 1^st^ cleavage.

## Acknowledgment

We thank R. Haley for helpful comments. We thank A. Fire for the gift of plasmids pPD95.69 and pPD117.01. We thank CGC, which is funded by NIH Office of Research Infrastructure Programs (P40 OD010440), for providing some strains.

## Author Contributions

The author(s) have made the following declarations about their contributions: Conceived and designed the experiments: YF ZZ. Performed the experiments: YF OPR ZL ZZ. Analyzed the data: YF OPR ZZ. Prepare the manuscript: YF OPR ZZ.

## Supplemental figure legends

**Figure S1. The relative Ca^2+^ signal intensity over time in live or necrotic PLM neurons in different genotypes and/or environmental conditions**

*(Related to Figures 2 and 4)*

Presented here are results of time-lapse recording experiments that monitor the intensity of cytoplasmic Ca^2+^ and cell swelling of PLM neurons during embryonic development. All embryos carry the transgenic array expressing P_*mec-7*_ *GCaMP5G*.

(A) DIC and fluorescence time-lapse images of two live PLM neurons in one *mec-4(+)* embryo. Time points are marked as min post-1^st^ embryonic division. White and yellow arrowheads mark the PLML and PLMR neurons, respectively. Scale bars are 10*µ*M.

(B) The relative signal levels of GCaMP5G were measured in 4 live PLM neurons and plotted over time. Graph (a) displays the plots of the PLML and PLMR neurons shown in (A), whereas graph (b) displays the plots of the PLML and PLMR neurons in an additional embryo.

(C) (a-b) The relative signal levels of GCaMP5G were measured in two necrotic PLM neurons in a *mec-4(e1611)* mutant embryo from a plate treated with DMSO but no dantrolene. Two other examples are shown in Figure 7. (c) The mean integral cytoplasmic Ca^2+^ intensities from multiple necrotic PLM neurons in *mec-4(e1611)* embryos and live PLM neurons in *mec-4(+)* embryos were obtained by calculating the integral value of GCaMP5G intensities within the time period of 560-800 min post-1^st^ embryonic division. Error bars indicate s.e.m. The numbers in paratheses in each bar represent the number of PLM neurons quantified.

(D) (a-b) The relative GCaMP5G signal levels measured in 2 necrotic PLM neurons in *mec-4(e1611)* mutant embryos from a 3μM dantrolene treated plate are plotted over time.

**Figure S2. NH_4_Cl partially suppresses necrosis but allows the exposure of PS on live PLM neurons**

*(Related to Figures 7)*

(A) The NH_4_Cl treatment partially suppresses necrosis induced by the *mec-4(e1611)* mutation. The mean numbers of PLM neurons (labeled with P_*mec-7*_GFP) that display the swelling necrosis phenotype in the tails of young *ced-1(e1735); mec-4(e1611)* L1 larvae from liquid cultures treated or not treated with 5mM NH_4_Cl are presented in the graph. Bars represent the mean values of each sample. Error bars indicate s.e.m. The numbers in the parentheses represent the numbers of L1 larvae scored. “**”, 0.001<p<0.01, Student *t*-test.

(B) Images of necrotic and live PLM neurons that present PS on their outer surfaces in *mec-4(e1611)* L1 larvae. L1 larvae were from liquid cultures either treated with 5mM NH_4_Cl (d-f, white arrowheads) or left untreated (a-c). PLM neurons (c (white arrow), f (white and yellow arrowheads)) are labeled with the P_*mec-7*_GFP reporter. DIC images identify one necrotic PLM neuron (white arrow) and one apoptotic cell (yellow arrow) in the tail of an L1 larva not treated with NH_4_Cl (a), and one necrotic PLM (yellow arrowhead) and one live PLM (white arrowhead) in an L1 larva from the 5mM NH_4_Cl treated culture (d-f). PS presentation on the cell surfaces (b, e) is detected by MFG-E8::mCherry. Scale bars are 5*µ*m.

