## Supplementary figures and images for "Calcium ions trigger the exposure of phosphatidylserine on the surface of necrotic cells"

### Supplemental Figure 1

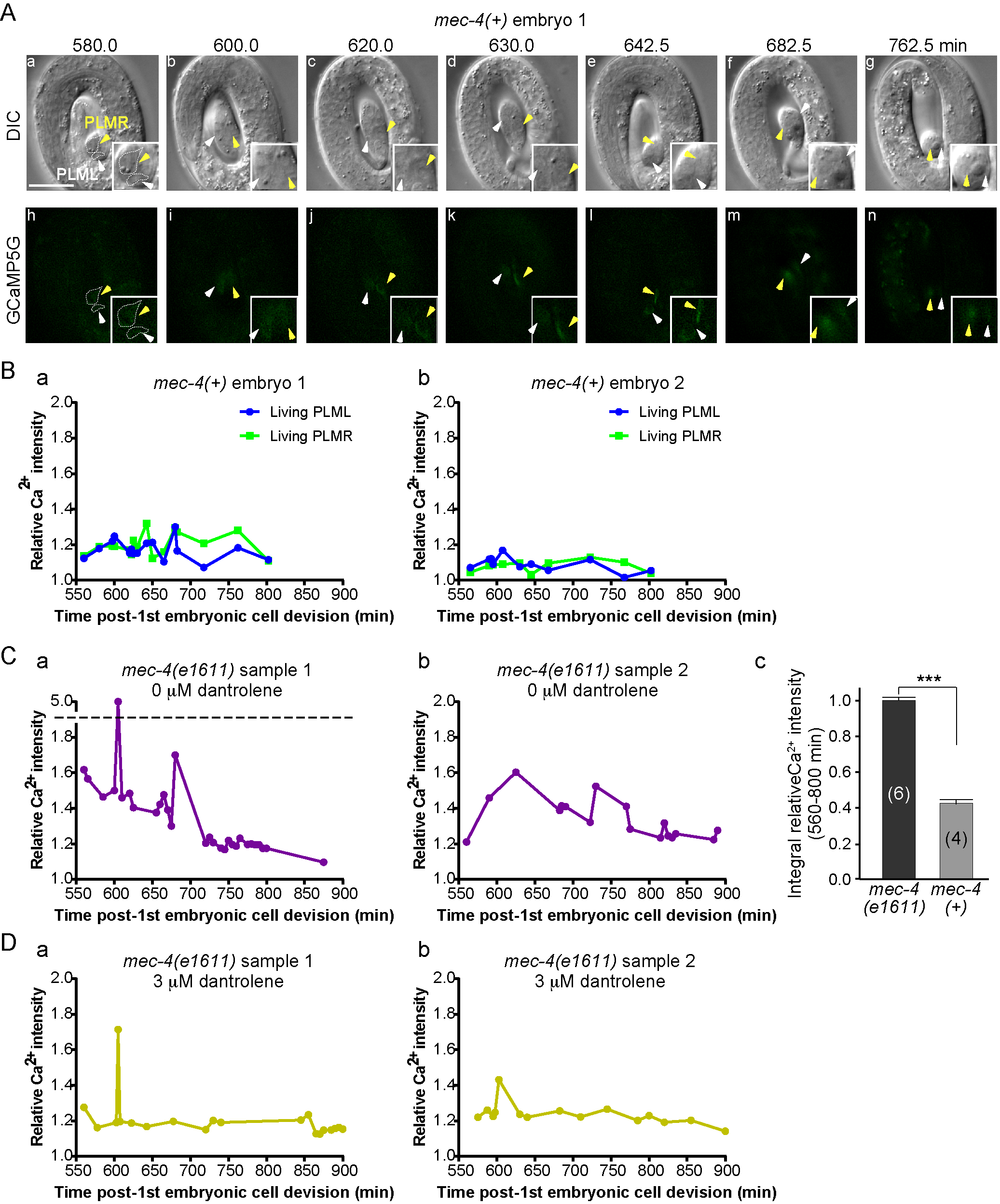

### Supplemental Figure 2

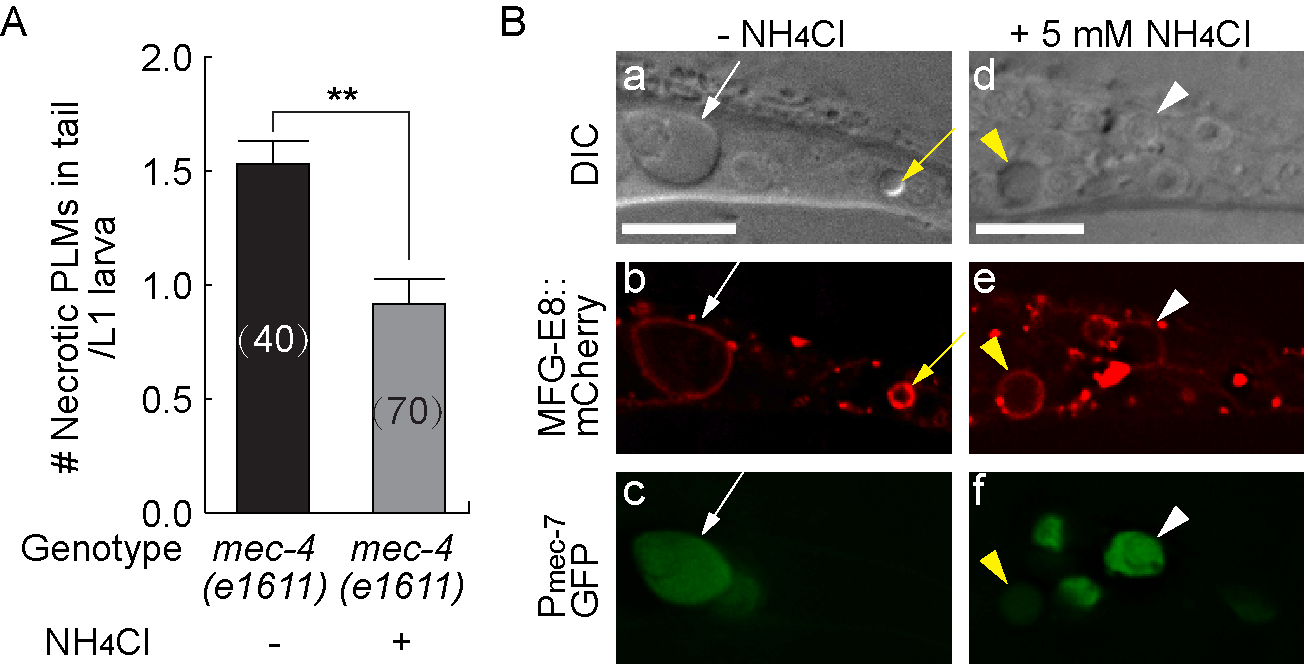
